# *In-vitro* reconstitution of Herpes Simplex Virus 1 fusion identifies low pH as a fusion co-trigger

**DOI:** 10.1101/2023.09.08.556861

**Authors:** J. Martin Ramirez, Ariana Calderon-Zavala, Ariane Balaram, Ekaterina E. Heldwein

**Affiliations:** Department of Molecular Biology and Microbiology, Tufts University School of Medicine, Boston, MA 02111; Graduate Program in Molecular Microbiology, Graduate School of Biomedical Sciences, Tufts University School of Medicine, Boston, MA 02111; Medical Scientist Training Program, Graduate School of Biomedical Sciences, Tufts University School of Medicine, Boston, MA, 02111

**Keywords:** HSV-1, VSV pseudotype, membrane fusion, *in-vitro* fusion, lipids, synthetic liposomes, HVEM, nectin-1, receptor, glycoproteins, low pH, trigger

## Abstract

Membrane fusion mediated by Herpes Simplex virus 1 (HSV-1) is a complex, multi-protein process that is receptor-triggered and can occur both at the cell surface and in endosomes. To deconvolute this complexity, we reconstituted HSV-1 fusion with synthetic lipid vesicles *in vitro*. Using this simplified, controllable system, we discovered that HSV-1 fusion required not only a cognate host receptor but also low pH. On the target membrane side, efficient fusion required cholesterol, negatively charged lipids found in the endosomal membranes, and an optimal balance of lipid order and disorder. On the virion side, the four HSV-1 entry glycoproteins gB, gD, gH, and gL were sufficient for fusion. We propose that low pH is a biologically relevant co-trigger for HSV-1 fusion. The dependence of fusion on low pH and endosomal lipids could explain why HSV-1 enters most cell types by endocytosis. We hypothesize that under neutral pH conditions, other, yet undefined, cellular factors may serve as fusion co-triggers. The *in-vitro* fusion system established here can be employed to systematically investigate HSV-1-mediated membrane fusion.

**IMPORTANCE:** Herpes simplex virus 1 (HSV-1) causes life-long, incurable infections and diseases ranging from mucocutaneous lesions to fatal encephalitis. Fusion of viral and host membranes is a critical step in HSV-1 infection of target cells that requires multiple factors on both the viral and host sides. Due to this complexity, many fundamental questions remain unanswered, such as the viral and host factors that are necessary and sufficient for HSV-1-mediated membrane fusion and the nature of the fusion trigger. Here, we developed a simplified *in-vitro* fusion assay to examine the fusion requirements and identified low pH as a co-trigger for virus-mediated fusion *in vitro.* We hypothesize that low pH has a critical role in cell entry and, potentially, pathogenesis.

## INTRODUCTION

Enveloped viruses must fuse their membranes with host cell membranes to initiate infection. This process has a high kinetic barrier that is overcome by viral surface proteins called membrane fusogens (1). Membrane fusogens aid in overcoming the energetic barrier by forming an extended structure that inserts into the host membrane and folds back on itself, bringing the two membranes close enough together to fuse (2, 3). Fusogens are subject to temporospatial regulation that ensures their deployment only when certain conditions are met, such as in the presence of specific host cell factors, target membrane lipid composition, or environmental triggers such as low pH of internal cellular compartments, or a combination thereof (4, 5). Depending on the target host cell and the virus, said triggers determine when and where the viral envelope will fuse with the host cell membrane to deliver the viral genome into the target host cell.

Herpesviruses are double-stranded-DNA enveloped viruses that are endemic and establish lifelong latent, reactivating infections (6). The prototypical herpes simplex virus 1 (HSV-1) initially infects epithelial cells in the mucosa or the skin and uses retrograde transport in axons to establish latency in the dorsal root ganglia (7). HSV-1 is found in at least two-thirds of the world’s population (6). In addition to oral and genital lesions, HSV-1 can cause keratitis – a major cause of blindness worldwide, and temporal lobe encephalitis (8, 9).

Herpesviruses penetrate target cells by fusing their lipid envelopes with a host cell membrane. Their mechanisms for entry into the host cell are some of the most complex among viruses because they employ a large number of viral glycoproteins and host receptors. In HSV-1, entry-associated functions are distributed across four viral glycoproteins: gD, gH, gL, and gB [reviewed in (10–12)]. According to the current model, these four viral glycoproteins orchestrate membrane fusion through a sequential activation process termed regulatory cascade (13) [reviewed in (10, 14, 15)]. First, the receptor-binding glycoprotein, gD, binds one of its three cognate receptors on the surface of the host cell and undergoes a conformational change, enabling it to bind and activate the gH/gL complex. The gH/gL complex, in turn, transmits the activating signal from the receptor-bound gD to gB, a viral fusogen that mediates the merger of the viral and host membranes. To add to this complexity, the HSV-1 envelope contains twelve other glycosylated and unglycosylated surface proteins (16) that could influence cell entry [reviewed in (17)].

On the host side, HSV-1 engages a variety of host molecules for attachment, internalization, and other entry-related functions, in a cell-specific manner [reviewed in (17)]. Entry requires one of three entry receptors: nectin-1, a member of the immunoglobulin superfamily that functions as a cell adhesion molecule; the herpesvirus entry mediator (HVEM), a member of the tumor necrosis factor receptor superfamily; or a 3-O-sulfonated derivative of heparan sulfate (3-OS-HS) [reviewed in (18)]. These three receptors are mutually independent and do not function merely as attachment factors. Instead, they trigger a cascade of conformational changes in the viral glycoprotein machinery described above. HSV-1 then utilizes distinct cell-type-dependent entry pathways, all of which require fusion of the viral envelope with either the host plasma membrane (19–21), the membrane of an endocytic vesicle, or an endosome after internalization (22–27).

But aside from the host receptors, the fusion requirements on the host side – proteins, lipids, ions – have not yet been fully defined. Moreover, the question of what triggers fusion has not yet been settled. HSV-1-mediated fusion is commonly thought to be triggered by binding to one of the three host entry receptors (receptor-triggered) rather than by exposure to the low pH of the endosomal compartment (pH-triggered). This is because HSV-1 enters certain cell types, including neurons, by fusion at the plasma membrane, a process that occurs at neutral pH (19–21). Likewise, robust cell-cell fusion is observed at neutral pH in uninfected cells overexpressing HSV-1 gB, gH, gL, and gD in the presence of a cognate receptor (28, 29). However, HSV-1 enters some cell types, e.g., primary human keratinocytes, by endocytosis and fusion with an endosomal membrane (22–27), and entry is sensitive to inhibitors of endosomal acidification (30). Low pH triggers fusion in many viruses that use the endocytic route of entry. Therefore, low pH could, in principle, act as a potential fusion trigger in HSV-1.

Our current mechanistic models of HSV-1 membrane fusion are primarily based on studies of cell-cell fusion of uninfected cells overexpressing the four HSV-1 entry glycoproteins. However, the cell-cell fusion system, while informative, does not adequately recapitulate the fusion of viral particles with target membranes due to differences in the geometry of the opposing membranes, membrane composition, and possible contributions by the other viral glycoproteins. Studying membrane fusion in the context of HSV-1 infection provides a more authentic system. However, such studies are confounded by the complex, dynamic nature of the host cell and do not measure fusion directly because they utilize downstream reporters. Therefore, studying HSV-1-mediated membrane fusion requires an experimentally tractable system that recapitulates the important aspects of viral fusion and can be easily manipulated.

Here, we reconstituted the HSV-1 fusion process for the first time *in vitro* with purified virions and synthetic lipid vesicles of defined composition. Using this simplified, controllable system, we discovered that HSV-1 fusion required not only a cognate host receptor but also low pH. We also found that on the target membrane side, efficient fusion required cholesterol, negatively charged lipids found in the endosomal membranes, and an optimal balance of lipid order and disorder. Finally, by using the Vesicular Stomatitis Virus pseudotypes, we confirmed that on the virion side, the four HSV-1 entry glycoproteins gB, gD, gH, and gL were sufficient for fusion.

We hypothesize that low pH is a biologically relevant fusion co-trigger for HSV-1. The dependence of fusion on low pH and endosomal lipids could explain why HSV-1 chooses the endocytic route of entry into most cell types. Membrane fusion under neutral pH conditions – in the context of cell entry at the plasma membrane or cell-cell fusion – may use other, yet undefined, cellular factors as co-triggers. The *in-vitro* fusion system established here can be used for further systematic exploration of HSV-1-mediated fusion.

## RESULTS

### The in-vitro fusion setup

Virion fusion with artificial membranes (lipid vesicles or lipid bilayers) has been reconstituted for several viruses (31–43). In a typical *in-vitro* fusion experiment, virions or liposomes are labeled with either a lipophilic or an aqueous fluorescent dye. Changes in fluorescence upon lipid mixing or content mixing report on membrane hemifusion or full fusion, respectively. These can be due to the dequenching of a dye incorporated at self-quenching concentrations, an increase or a decrease in FRET efficiency, or changes in the fluorescence emission profile. Triggers of membrane fusion vary depending on the virus and can include receptors, co-receptors, or environmental triggers such as pH or ions.

In our experiments, we chose to monitor content mixing, i.e., the mixing of the viral interior with that of the liposome, which reports on the full fusion of the opposing membranes. This was done by incorporating an aqueous fluorophore, Sulforhodamine B (SRB), into the target liposomes at self-quenching concentrations (**Fig. 1A**). Membrane fusion leads to mixing of the viral and liposome interiors, which dilutes the aqueous fluorophore, causing it to de-quench and emit higher fluorescence. An increase in fluorescence reports on content mixing (**Fig. 1B**). SRB was chosen as a reporter because it cannot traverse membranes and is not pH sensitive, which allows monitoring of pH-triggered membrane fusion. It has been successfully used to monitor *in-vitro* fusion for several viruses (35, 37).

**Figure 1.**
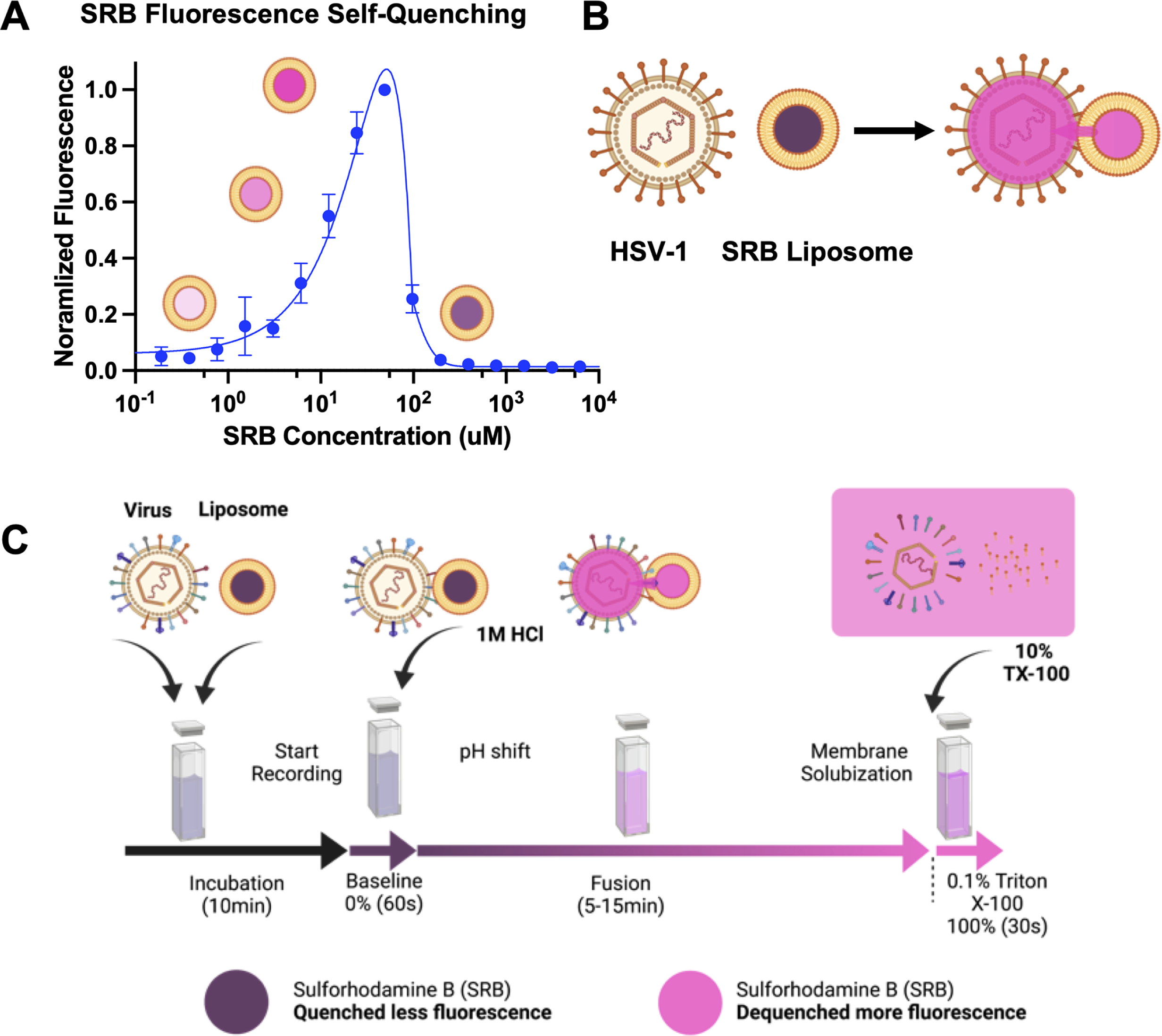
The *in-vitro* SRB dequenching assay measures content mixing and fusion. (A) Sulforhodamine B (SRB) self-quenches at mM concentrations. Serial dilutions of SRB demonstrate the self-quenching capacity of SRB as concentration increases. (B) A schematic representation of content mixing between an enveloped virus and a liposome loaded with SRB at a self-quenching concentration. When fusion occurs, the viral and the liposome contents mix, and the SRB is diluted and de-quenched, which increases its fluorescence. (C) Bulk fusion workflow. Virions and liposomes are incubated for 10 min (black arrow) before the start of fluorescence recording. Baseline fluorescence is recorded for 1 min (purple arrow), at which point the reaction is triggered by pH shift. Fluorescence is recorded for an additional 5-15 minutes (purple to pink arrow) before the addition of 10% Triton X-100 in HBS-citrate buffer to a final concentration of 0.1%. Triton X-100 causes full SRB dequenching, which is set to 100% for normalization. Illustrations of the process are shown above the arrows. Diagrams and cartoons were created using BioRender.com.

To optimize the experimental setup, we used the Vesicular Stomatitis virus (VSV), for which the fusion requirements are well defined (44, 45) and the *in-vitro* fusion conditions have been established (42, 46-48). VSV-mediated fusion is triggered by acidification (49–52). An earlier study (42) that measured hemifusion by monitoring FRET between two lipophilic dyes incorporated into target liposomes reported that VSV-mediated fusion *in-vitro* required 1-palmitoyl-2-oleoyl-sn-glycero-3-phospho-L-serine (POPS) in target membranes, which may act as a receptor (42). Therefore, we used large unilamellar vesicles (LUVs) composed of 33% 1-palmitoyl-2-oleoyl-glycero-3-phosphocholine (POPC), 16.7% 1-palmitoyl-2-oleoyl-sn-glycero-3-phosphoethanolamine (POPE), 16.7% POPS, and 33% cholesterol (POPC:POPE:POPS:Chol = 2:1:1:2) (**Table 1: POPS**) (42). In the same study, optimal fusion under *in-vitro* conditions was observed at pH values <5.6 (42), so we tested a pH range between 5.0 and 7.4. Lastly, to ensure homogeneity, SRB-containing liposomes were evaluated for size and concentration by static light scattering (**Supplementary Fig. S1A-C).**

**Table 1.** Lipid compositions of LUVs used in this work. **POPS** (palmitoyl-2-oleoyl-sn-glycero-3-phospho-L-serine), **BMP** (bis(monooleoylglycero)phosphate), **POPA** (palmitoyl-2-oleoyl-sn-glycero-3-phosphate), **POPE** (palmitoyl-2-oleoyl-sn-glycero-3-phosphoethanolamine), **Chol** (Cholesterol), **SM** (sphingomyelin), **DGS-NTA (Ni**) (1,2-dioleoyl-sn-glycero-3-[(N-(5-amino-1-carboxypentyl)iminodiacetic acid)succinyl] (nickel salt).

Purified VSV virions were mixed with SRB-labeled large unilamellar vesicles (LUVs) in a fluorescence cuvette, and the emission intensity of SRB was recorded as a function of time (**Fig. 1C and Supplementary Fig. 2A**). Since absolute fluorescence emission values can vary across liposome preparations, fusion values across different experiments were normalized. The initial fluorescence emission intensity after virion and liposome mixing, but before triggering, was defined as 0% fusion (**Fig. 1C and Supplementary Fig. 2A**). The detergent Triton X-100 was added at the end of each experiment to disperse the membranes and measure the fully unquenched fluorescence emission intensity. This value was defined as 100% fusion. In each experiment, full fusion, i.e., content mixing, was assessed by an increase in the emission intensity due to SRB dequenching and normalized to fully unquenched fluorescence emission intensity (**Supplementary Fig. 2A**) by using the equation: *F_normalized_=F_t_-F_i_ / (F_max_-F_i_)^−1^ x 100%*, where F_t_ is fluorescence at time *t, F_i_* is initial fluorescence and *F_max_*is fluorescence after treatment with Triton X-100.

To compare multiple data sets, the normalized values at the endpoint of each experiment, which correspond to values just before the addition of Triton X-100, were averaged and plotted as bar graphs **(Supplementary Fig. 2A**). As expected, significant fluorescence emission dequenching, ∼40-45%, indicative of fusion, was observed at pH values ≤5.5 **(Supplementary Fig. 2A and 2B**). At neutral pH, the fluorescence signal was much lower. These results were in agreement with a previous study (42) despite the use of a different fluorescent reporter, SRB, which reports on fusion instead of hemifusion and uses LUVs as opposed to small unilamellar vesicles (SUVs, r<50nm).

### HSV-1 fusion in vitro requires both a receptor and low pH

Having validated the *in-vitro* bulk fusion assay with VSV, we next moved on to HSV-1. HSV-1 was propagated and titrated on Vero cells. To ensure batch-to-batch consistency, HSV-1 virions were purified by density gradient centrifugation. HSV-1-infected cells typically produce not only virions but also particles lacking capsids, which are known as light particles (L-particles). Since L-particles have a different glycoprotein composition (53) and may have different fusogenic requirements from the virions, HSV-1 virions were separated from L-particles by density gradient centrifugation using a 10-50% iodixanol gradient (**Supplementary Fig. S1D**). The purified HSV-1 virions were analyzed and quantified using multi-angle light scattering (MALS) and dynamic light scattering (DLS) (**Supplementary Fig. S1E-H).** The typical particle radius was ∼98 nm (**Supplementary Fig. S1G**) and agreed with the previously reported range of 92-115 nm (54, 55). Particle counts (**Supplementary Fig. S1E**) were used to calculate the particle-per-PFU ratios (**Supplementary Fig. S1H**). Only the virion preparations with the particle-to-PFU ratio of 20-30 were deemed to be of sufficiently high quality (56).

To measure HSV-1-mediated fusion, purified HSV-1 virions were incubated with SRB-labeled LUVs in a fluorescence cuvette. Initially, we used liposomes of the same composition as in the VSV-mediated fusion experiment, 33% POPC, 16.7% POPE, 16.7% POPS, and 33% cholesterol (POPC:POPE:POPS:Chol = 2:1:1:2) (**Table 1: POPS**). Since HSV-1-mediated fusion is receptor-triggered, we used the purified recombinant soluble ectodomain of HVEM, HVEM200t-His_8_ to trigger fusion (**Supplementary Fig. S3A).** HVEM200t-His_8_ was added to the HSV-1/LUV mixture after incubation. However, at neutral pH, no significant SRB dequenching was observed (**Fig. 2A and D**). Since HSV-1 enters most known target cell types by pH-dependent endocytosis (10, 11, 57, 58), we tested pH ranging from pH 6.5 to 4.5 by adding predetermined amounts of HCl after incubation with a soluble receptor. Upon acidification in the presence of HVEM200t-His_8_, we observed robust fusion at pH <5.5, ∼40% (**Fig. 2A and D**).

**Figure 2.**
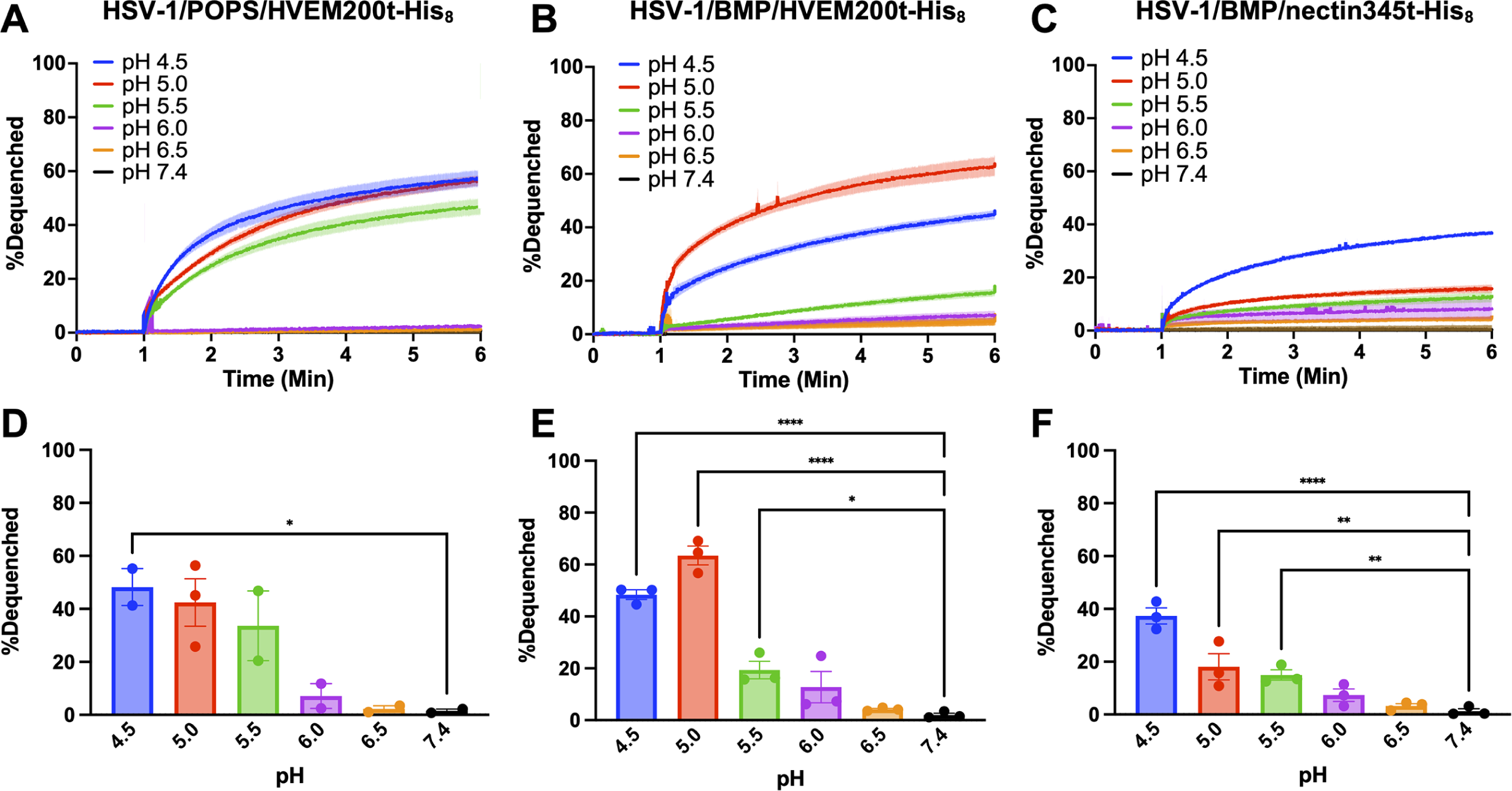
A soluble receptor (HVEM or nectin-1) in combination with low pH triggers HSV-1 fusion *in vitro*. HSV-1 was incubated with SRB-containing LUVs of different lipid compositions at different pH in the presence of HVEM200t-His_8_ or nectin345t-His_8_. Increase in SRB fluorescence due to its dilution and dequenching reported on fusion. Lipid compositions are listed in Table 1. Fluorescence was recorded and normalized to fully de-quenched fluorescence by using the equation: *F_normalized_=F_t_-F_i_ / (F_max_-F_i_)^−1^ x 100%*, where *F_t_*is fluorescence at time *t, F_i_* is initial fluorescence and *F_max_* is fluorescence after the addition of Triton X-100. (A-C) For each condition, representative traces of single biological replicates, each consisting of three technical replicates, are shown. Curves represent the mean values, and the shaded area, the SEM. (D-F) The extent of fusion at 5 min post-triggering (acidification). Each data point is a biological replicate representing a mean normalized fluorescence from three technical replicates. Bars represent the mean values, and the error bars, the SEM. Each condition was done in triplicate except for HSV-1/POPS/HVEM200t-His_8_, which was done in duplicate. *: p < 0.05, **: p < 0.01, ***: p<0.001 ****: p<0.0001.

Although POPS worked consistently in the *in-vitro* fusion experiments with VSV **(Fig. 1D and E**), with HSV-1, we observed variability in fluorescence at intermediate pH values such that statistical significance was only achieved at pH 4.5 (**Fig. 2D),** and biological replicates did not have the same trend (**Supplementary Fig. S4A**). Therefore, we replaced POPS with another lipid with a negatively charged headgroup, bis(monooleoylglycero)phosphate (BMP), which is specific to late endosomes. The resulting liposomes contained 33% POPC, 16.7% POPE, 16.7% BMP, and 33% cholesterol, (POPC:POPE:BMP:Chol = 2:1:1:2) (**Table 1: BMP**). With the BMP liposomes, robust fusion was observed at pH ≤5.0 (**Fig. 2B and E**), just as with the POPS liposomes (**Fig. 2A and D**). However, in the presence of BMP, the fusion increased to 60%, and reproducibility improved, with fusion reaching statistical significance at pH values ≤5.5 (**Fig. 2E**). Individual biological replicates had very similar trends (**Supplementary Fig. S4B**).

To assess the potential contributions of liposome leakage, liposome rupture, or mechanical perturbation due to stirring to the fluorescence signal, and to determine the optimal end-point time for measurements, we tested the BMP liposomes (**Table 1: BMP**) under conditions non-permissive for fusion by leaving out one or more required components, e.g., HSV-1, HVEM200t-His_8_, or low pH (**Supplementary Fig. S5A**). With LUVs alone at pH 7.4, the final dequenching values were below the baseline indicating moderate photobleaching instead of fluorescence (**Supplementary Fig. S5, brown**). The addition of HVEM200t-His_8_ or HSV-1 to the LUVs at pH 7.4 minimally increased the fluorescence signal (**Supplementary Fig. S5, orange and black, respectively**) to <10% after an hour. However, with LUVs at pH 5.0, either alone or in the presence of HVEM200t-His_8_, we detected a time-dependent increase in fluorescence (**Supplementary Fig. S5, green and red, respectively**) ∼40%, which indicated that some liposome leakage or rupture was occurring at low pH. Under these conditions, the fluorescent signal of ∼20% was the lowest at 5 min post-triggering (**Supplementary Fig. S5D, green and red**). Importantly, under conditions enabling fusion, i.e., at pH 5.0 and in the presence of both HSV-1 and HVEM200t-His_8_, the fluorescence signal nearly plateaued at 5 min post-triggering, reaching ∼60% (**Supplementary Fig. S5D, blue**). Therefore, to minimize any potential contributions of liposome leakage to the fluorescence signal, we chose 5 min post-triggering as the endpoint for all subsequent *in-vitro* fusion measurements. Surprisingly, with LUVs at pH 5.0 in the presence of HSV-1 (without HVEM200t-His_8_), the fluorescence signal was minimal (**Supplementary Fig. S5, purple**) with <10% fluorescence. This suggested that liposome leakage caused by low pH was somehow mitigated by the presence of HSV-1, effectively ruling out nonspecific liposome leakage as a significant contributor to the fusion signal under fusion-permitting conditions.

We next tested fusion in the presence of the soluble ectodomain of another HSV-1 receptor, nectin-1, nectin345t-His_8_ (**Supplementary Fig. S3B**). Just as with HVEM200t-His_8_, significant fusion was observed only upon acidification, with fusion extent reaching ∼40% at pH 4.5 (**Fig. 2C and F**) and having a consistent trend among biological replicates (**Supplementary Fig. S4C**). In the presence of nectin345t-His_8_, fusion was somewhat less efficient, ∼40%, compared to ∼60% observed in the presence of HVEM200t-His_8_. Additionally, in the presence of nectin345t-His_8_ the highest fusion was observed at pH 4.5 instead of pH 5.0. Despite these differences, HSV-1-mediated fusion *in vitro* in the presence of either HVEM200t-His_8_ or nectin345t-His_8_ required low pH.

The differences in fusion extent between the two receptors could be due to the differences in fusion rates or fusion efficiency. To resolve these alternate possibilities, we calculated the fusion rate constants for fusion in the presence of HVEM200t-His_8_ or nectin345t-His_8_ at pH 4.5 or 5.0 by fitting the normalized data using analyses built into GraphPad PRISM9 (**Supplementary Fig. 7**). Single exponential functions did not fit the data, so the two-phase (exponential) association was used instead, which yielded two rate constants, k_fast_ and k_slow_. Of the two, k_fast_ is the more relevant parameter because it describes the rate of the fast exponential component, which likely specifically represents the fusion process, whereas k_slow_ could represent the rate of a slow background process or the nonspecific dequenching observed in negative controls. k_fast_ was relatively similar across the conditions (**Supplementary Fig. 7A**). This suggests that the fusion mechanism is the same regardless of the receptor, which is consistent with the current model [reviewed in (10, 14, 15)]. Therefore, we hypothesize that the differences in fusion extent in the presence of HVEM200t-His_8_ or nectin345t-His_8_ are due to differences in fusion efficiency.

### Incubation of HSV-1 with a soluble receptor and low pH reduces its infectivity

Fusion triggers activate viral fusogens by inducing a conformational change that causes the active, prefusion form to refold into the inactive, postfusion form (1). If the fusogen is triggered in the absence of a target membrane, it will refold into the postfusion form unproductively, which would effectively inactivate it. Therefore, if receptor and low pH functioned as co-triggers of the HSV-1 fusogenic machinery, then pre-treatment of HSV-1 with both co-triggers in the absence of the target membrane would inactivate it, thereby reducing the viral titer. Indeed, pre-treatment of HSV-1 with HVEM200t-His_8_ and low pH reduced its infectivity significantly, in a pH-dependent manner (**Fig. 3**), HVEM200t-His_8_ was tested because it had the most robust dequenching and would be used for other experiments. In contrast, pre-treatment of HSV-1 with HVEM200t-His_8_ at pH 7.4 or in the absence of receptor at pH 5.0, had a minimal, statistically insignificant effect on infectivity (**Fig. 3**). While the optimal pH for HSV-1 inactivation, pH 4.5 (**Fig. 3**), was lower than the optimal pH for *in-vitro* fusion, pH 5.0 (**Fig. 2D**), this discrepancy could be due to the differences between the two experimental setups. In the infection experiment, HSV-1 was incubated with HVEM200t-His_8_ at low pH for 1 hour in the absence of target cells. By contrast, in the *in-vitro* fusion experiment, HSV-1 was incubated with HVEM200t-His_8_ and target LUVs for 10 min before acidification, at which point the reaction was triggered by shifting the pH and monitored for 5 minutes. Regardless, both HSV-1 inactivation and *in-vitro* fusion require the presence of both HVEM200t-His_8_ and low pH.

**Figure 3.**
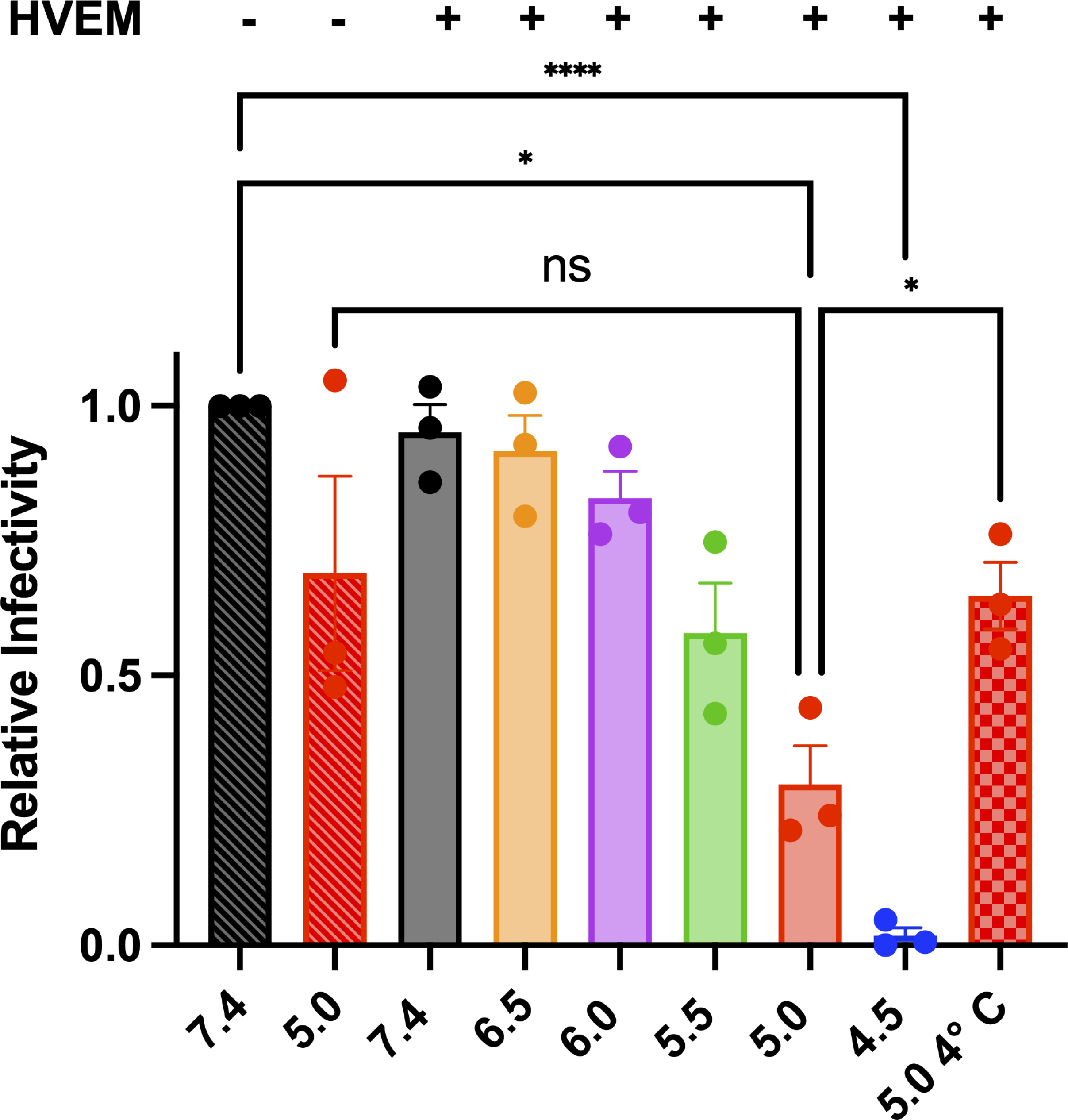
Pre-treatment with low pH and HVEM 200t-His_8_ inactivates HSV-1. HSV-1 was incubated with soluble HVEM200t-His_8_ for 1 hour and pH shifted at 37 °C or 4 °C for 1 hour. Reactions were back titrated to pH 7.4, diluted in minimal media, and titrated on Vero cells. PFUs were normalized to the no HVEM pH 7.4 condition (black crosshatched bar). Each data point represents a biological replicate. Each condition was done in triplicate. Bars represent the mean values, and the error bars, the SEM. *: p < 0.05, **: p < 0.01, ***: p<0.001 ****: p<0.0001.

### In-vitro HSV-1-mediated fusion requires a lipid with a negatively charged headgroup and cholesterol in the target membrane

To dissect the lipid requirements for HSV-1-mediated fusion *in vitro*, we systematically eliminated POPE, BMP, and cholesterol from target LUVs by replacing them with the equivalent amounts of POPC (**Table 1: no POPE, no BMP, no Chol**). Omission of BMP or cholesterol reduced fusion by ∼6-fold whereas omission of POPE did not (**Fig. 4**) Both BMP and POPS have negatively charged headgroups, and both supported HSV-1 fusion at similarly high levels, on average (**Fig. 2A, B, D, and E**). To test if other lipids with a negatively charged headgroup could support HSV-1 fusion, we tested 1-palmitoyl-2-oleoyl-sn-glycero-3-phosphate (POPA). POPA supported fusion at ∼2-fold lower level than BMP or POPS (**Fig. 4**). We conclude that HSV-1-mediated *in-vitro* fusion requires a lipid with a negatively charged headgroup on the target membrane side and that BMP and POPS are better than POPA.

**Figure 4.**
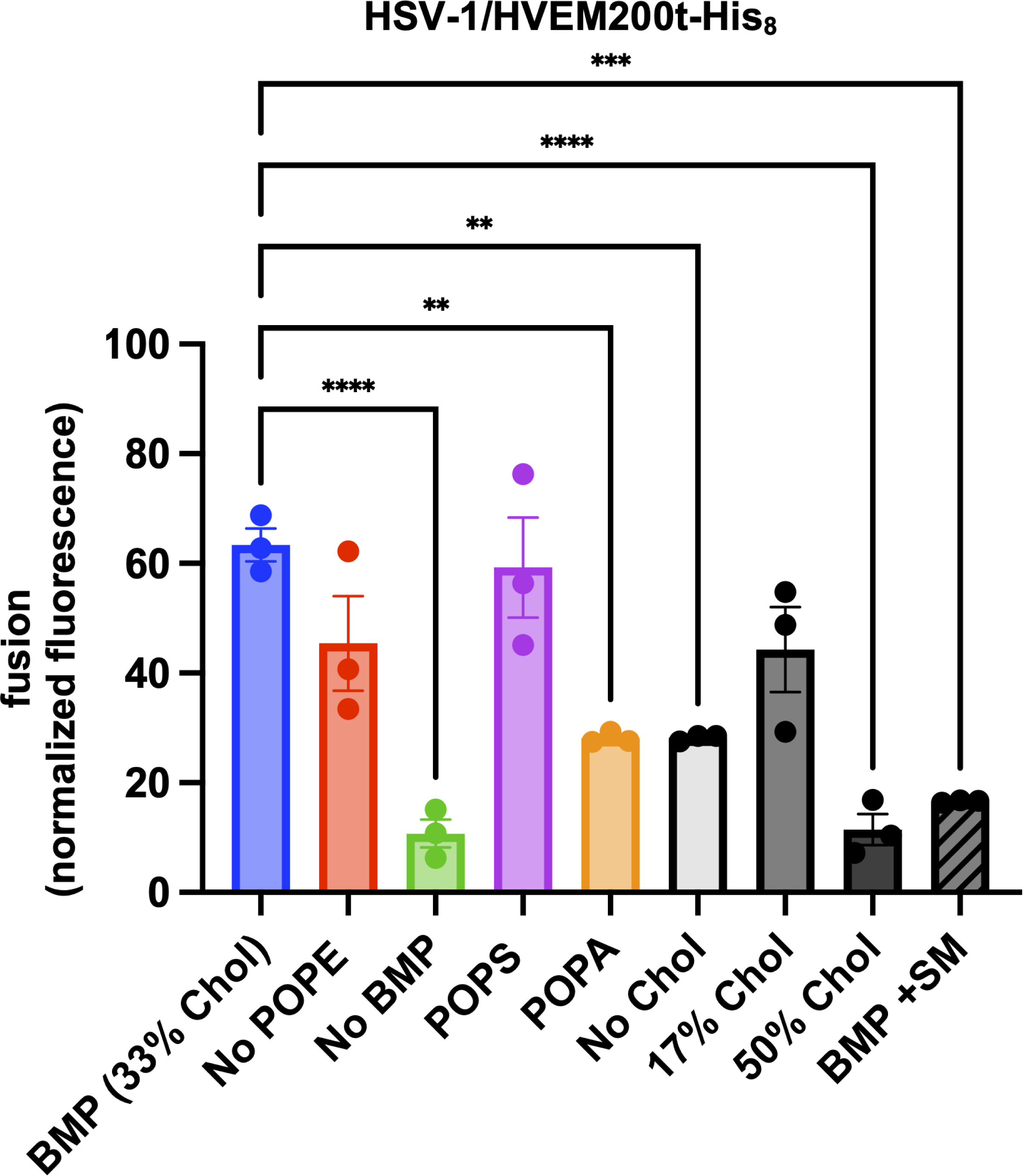
*In-vitro* fusion of HSV-1 depends on the lipid composition of the target membranes. HSV-1 was incubated with SRB-containing LUVs of different lipid compositions at pH 5.0 in the presence of HVEM200t-His_8_. Increase in SRB fluorescence due to its dilution and dequenching reported on fusion. Lipid compositions are listed in Table 1. Fluorescence was recorded at 5 min post-triggering (acidification) and normalized to fully de-quenched fluorescence by using the equation: *F_normalized_=F_t_-F_i_ / (F_max_-F_i_)^−1^ x 100%*, where *F_t_* is fluorescence at time *t, F_i_* is initial fluorescence and *F_max_* is fluorescence after the addition of Triton X-100. Each data point is a biological replicate representing a mean normalized fluorescence from three technical replicates. Each condition was done in triplicate. Bars represent the mean values, and the error bars, the SEM. Data for the BMP and POPS compositions are identical to those in Figure 2. Data for the BMP and POPS compositions are identical to those in Figure 2. *: p < 0.05, **: p < 0.01, ***: p<0.001 ****: p<0.0001.

Negatively charged lipids or cell surface glycosaminoglycans can act as initial attachment factors or receptors through electrostatic interactions (59, 60). We hypothesized that if BMP mediated HSV-1 attachment to the target membranes similarly, then displaying HVEM on the surface of the LUVs would bypass this requirement by allowing HSV-1 to attach to the LUVs by using gD/HVEM interaction. To achieve this, we turned to DGS-NTA, a Ni-chelating lipid that can be used, in conjunction with Ni, to anchor His-tagged proteins to liposomes (61, 62). We tested LUVs with or without BMP containing 3% DGS-NTA-Ni (**Table 1: BMP 3% DGS-NTA-(Ni), No BMP 3% DGS-NTA-(Ni**)). Unfortunately, we found that adding DGS-NTA-Ni to LUVs reduced fusion levels to <25% regardless of the BMP presence (**Supplementary Fig. S6**). Therefore, the role of the negatively charged lipid in HSV-1 fusion remained unclear.

Efficient *in-vitro* fusion also required cholesterol in the target LUVs (**Fig. 4**). Cholesterol is an important component of almost all cellular membranes and is required for the fusion of many enveloped viruses (40, 41, 63-70). We compared fusion in the presence of 0%, 17%, 33%, or 50% cholesterol (**Table 1: No Chol, 17% Chol, 33% Chol (BMP), 50% Chol**) and found 33% cholesterol to be the optimal amount (**Fig. 4**) producing ∼60% fluorescence. Fluorescence in the presence of 17% cholesterol was reduced to ∼40%, indicating reduced fusion levels, but this decrease was not statistically significant (**Fig. 4**).

### Efficient HSV-1-mediated fusion in vitro requires a balance of lipid order and fluidity

The levels of HSV-1-mediated fusion *in vitro* depended on the amount of cholesterol in the target membrane. 17 or 33% cholesterol supported efficient fusion at ∼40% and 60% fluorescence, respectively, whereas 0 or 50% cholesterol did not (**Fig. 4**), reducing fusion to ∼30% and ∼10%, respectively. In synthetic membranes, cholesterol has been shown to promote the formation of the liquid-ordered phase (L_o_), sometimes referred to as lipid rafts, that co-exists with the liquid-disordered phase (L_d_) (71–75). Therefore, we hypothesized that HSV-1-mediated fusion *in vitro* is influenced by membrane fluidity and lipid order.

To assess the relative amounts of L_o_ and L_d_ phases in our LUV preparations, we used the solvatochromic fluorescent dye 2-dimethylamino-6-lauroylnaphtalene (Laurdan). Laurdan can partition into L_d_ and L_o_ phases but changes its fluorescence emission maximum from 440 nm (L_o_) to 490 nm (L_d_), which makes it a useful reporter for L_o_/L_d_ transitions [reviewed in (76). Laurdan fluorescence emission in membranes is typically expressed as generalized polarization (GP) = (I_440nm_-I_480nm_)/(I_440nm_+I_480nm_)^−1^, where I is the fluorescence intensity at the indicated wavelength (actual peak values differ by the system but are usually within a few nm of 440nm and 480nm). Higher GP values indicate higher lipid order, i.e., L_o_>L_d_, whereas lower GP values indicate lower lipid order and higher fluidity, i.e., L_d_>L_o_.

As controls for membranes with high vs. low lipid order, we used LUVs composed of POPC, cholesterol, and sphingomyelin (SM) (POPC:Chol:SM=1:1:1) as well as LUVs composed of POPC and SM (POPC:SM=1:1) (**Table 1: PC:Chol:SM, PC:SM**). SM is a saturated lipid that is commonly found in biological lipid rafts (77, 78). Synthetic membrane bilayers composed of POPC:Chol:SM=1:1:1 form L_o_ phases (74) and are used to model lipid rafts. By contrast, synthetic membrane bilayers composed of POPC:SM=1:1 form L_d_ at temperatures above ∼ 40°C (the phase transition temperature of SM) and a combination of solid-ordered (S_o_) and L_d_ phases at temperatures below ∼40 °C (79).

In all compositions tested, 1% POPC was substituted with Laurdan. Initially, Laurdan emission spectra were recorded at 37 °C (**Fig. 5A**) and GP values were calculated (**Fig. 5B**). As expected, in POPC:Chol:SM LUVs, Laurdan had a strong 440-nm emission peak and GP>0, consistent with high amounts of L_o_ phase (**Fig. 5A and B**). Conversely, in POPC:SM LUVs, it had a strong 490-nm emission peak and GP<0, indicating predominantly the L_d_ phase (**Fig. 5A and B**). To characterize the thermal stability of the ordered lipid phases, we recorded Laurdan emission spectra in the 0-60 °C range and calculated the GP values (**Fig. 5C**). POPC:Chol:SM LUVs maintained high GP values at increasing temperatures, demonstrating the stability of L_o_ domains (**Fig. 5C, violet**). By contrast, the POPC:SM LUVs underwent a phase transition above 38 °C caused by the “melting” of the S_o_ domains, and the GP changes from positive to negative (**Fig. 5C, dark blue**).

**Figure 5.**
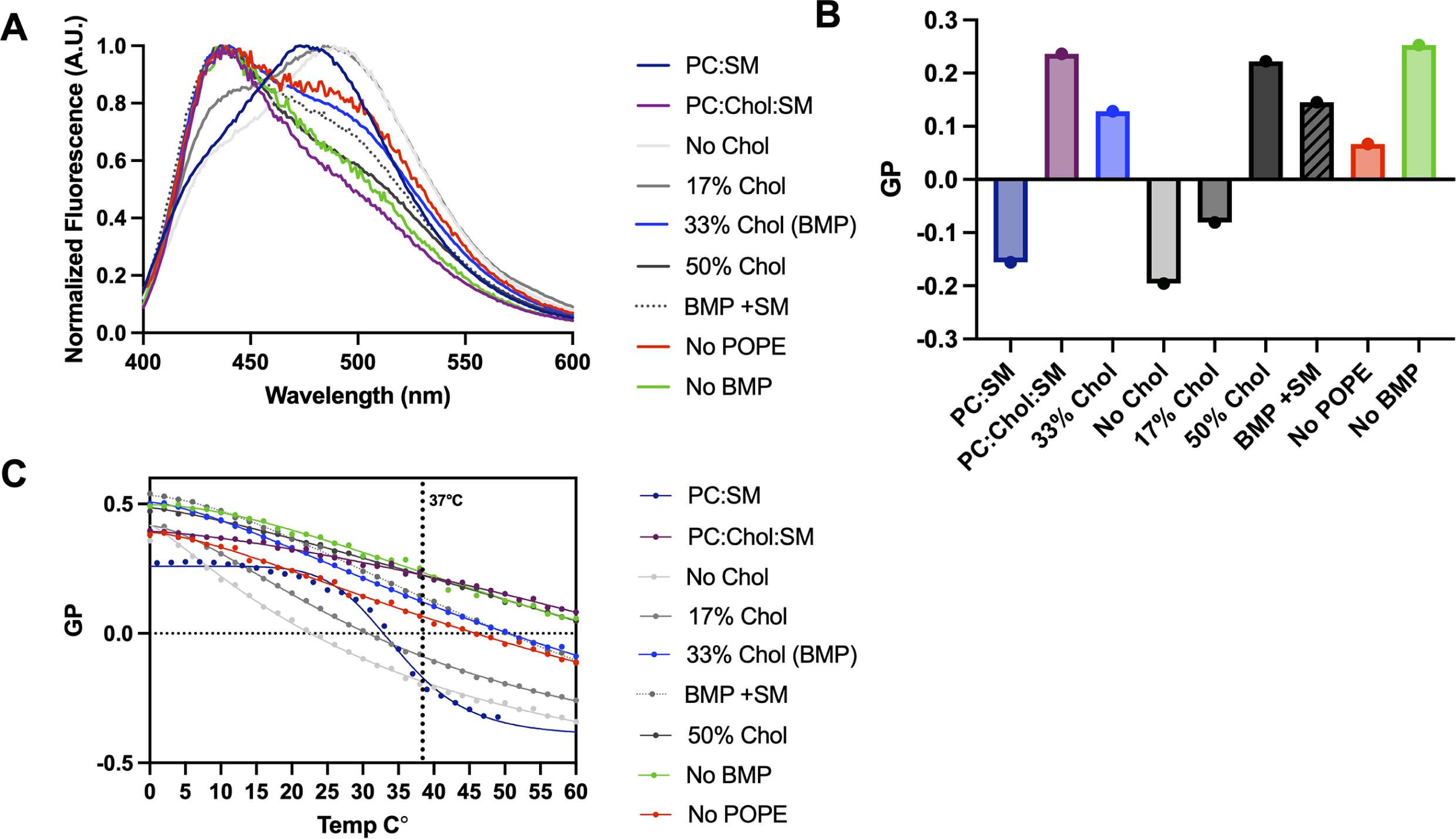
Estimates of lipid order of the synthetic membranes used in this study. (A) Laurdan fluorescence emission reports on lipid order composition. LUVs were prepared as for fusion experiments (Table 1) except that 1% POPC was substituted with Laurdan. Fluorescence emission was recorded and normalized from 400-600 nm. The emission peak at 440 nm reports the presence of an ordered lipid phase whereas the emission peak between 480 and 494 nm indicates a predominantly disordered lipid phase. PC:SM and PC:Chol:SM compositions served as controls for highly ordered and highly disordered phases, respectively. (B) Peak fluorescence values at 440 nm and 480 or 494 nm were used to calculate generalized polarization (GP) defined as I_440nm_-I_494nm_ / I_440nm+_I_494nm_, which reports on the extent of membrane order, with GP values increasing with increasing lipid order. (C) Fluorescence emission was recorded as a function of temperature, and GP values were calculated as in (B). For each condition, a single replicate was obtained.

Next, we tested LUVs used in the *in-vitro* fusion assays (**Table 1: No Chol, 17% Chol, 33% Chol (BMP), 50% Chol, No POPE, No BMP**). The GP values increased with the increasing cholesterol amount (**Fig. 5B**). The LUVs containing 33% or 50% cholesterol (**Table 1: 33% Chol (BMP), 50% Chol**) were relatively thermostable, indicating a higher order (**Fig. 5C, blue and black, respectively**), whereas those with 0% or 17% cholesterol (**Table 1: 0% Chol, 17% Chol**) were not, indicating higher fluidity (**Fig. 5C, light grey and dark grey, respectively**). These results are consistent with the known lipid ordering effect of cholesterol (80–83).

Interestingly, the GP and thermostability of No BMP LUVs were higher than BMP LUVs despite both having 33% cholesterol and on par with 50% Chol LUVs and POPC:Chol:SM LUVs (**Fig. 5B and C**). Conversely, the GP and thermostability of No POPE LUVs were lower than BMP LUVs, despite both having 33% cholesterol (**Fig. 5B and C**). Thus, we hypothesize that BMP and POPE have opposite effects on the ordered lipid phase, with BMP decreasing it and POPE increasing it.

To relate fusion efficiency to lipid order, we compared HSV-1-mediated fusion levels **(Fig. 4**) to the GP values across different lipid compositions **(Fig. 5**). Fusion was lowest without BMP or in the presence of 50% cholesterol (**Table 1: No BMP, 50% Chol**) **(Fig. 4**), which are lipid compositions with the some of the highest GP values, >0.2, comparable to the POPC:Chol:SM, indicative of a high amount of ordered lipid phase (**Fig. 5**). This suggests that high lipid order prevents membrane fusion. On the opposite end of the order spectrum, No Chol LUVs had the lowest GP value, ∼-0.2, which corresponds to predominantly L_d_ phase (**Fig. 5**), and supported intermediate levels of fusion **(Fig. 4**). However, efficient fusion was observed with 17% Chol, BMP, and No POPE LUVs, which are lipid compositions that had intermediate GP values, between ∼-0.1 and ∼0.1. The optimal fusion was observed with BMP LUVs **(Fig. 4**). Therefore, we hypothesize that efficient HSV-1-mediated fusion *in vitro* requires an optimal balance of ordered lipid phase and fluidity, achieved in the presence of 33% cholesterol, BMP, and POPE.

### Sphingomyelin reduces fusion

Biological membranes typically contain SM found in lipid rafts (77, 78). To test the role of SM in HSV-1-mediated fusion, we added 8% SM to BMP LUVs (**Table 1: BMP + SM**). 8% SM was chosen because an earlier subcellular fractionation study reported that late endosomes, which contain BMP, also contain approximately 8% SM (84, 85). Surprisingly, in the presence of SM, fusion levels dropped almost to the fusion levels of 50% Chol LUVs, with <20% fluorescence (**Fig. 4**) while the GP remained the same (**Fig. 5B and C**). We hypothesize that ordered lipid phases formed with vs. without SM are qualitatively different. While ordered lipid phases formed in 33% cholesterol, 16.7% BMP, and 16.7 % POPE – our optimal target LUV composition – support efficient HSV-1 *in-vitro* fusion, the addition of SM promotes the formation of the ordered lipid phase that is less conducive to fusion.

### HSV-1 entry glycoproteins gB, gH, gL, and gD are sufficient for in-vitro fusion

HSV-1 has as many as 16 envelope proteins, but only four – gB, gH, gL, and gD – are essential for viral entry and cell-cell fusion (13, 86, 87). To test whether these four glycoproteins were sufficient for *in-vitro* fusion, we generated VSV virions lacking the native fusogen G and pseudotyped with HSV-1 gB, gH, gL, and gD (VSVΔG-BHLD). Previously, we showed that these VSVΔG-BHLD pseudotypes entered two types of HSV-1-susceptible cells (57, 88). Here, we found that the VSVΔG-BHLD pseudotypes fused with the target BMP LUVs. As with HSV-1, the *in-vitro* fusion of the VSVΔG-BHLD pseudotypes required both the receptor (HVEM200t-His_8_ or nectin345t-His_8_) and low pH (**Fig. 6**). The fusion levels of VSVΔG-BHLD in the absence of the receptor were very low, ∼10% (**Supplementary Fig. S7**).

**Figure 6.**
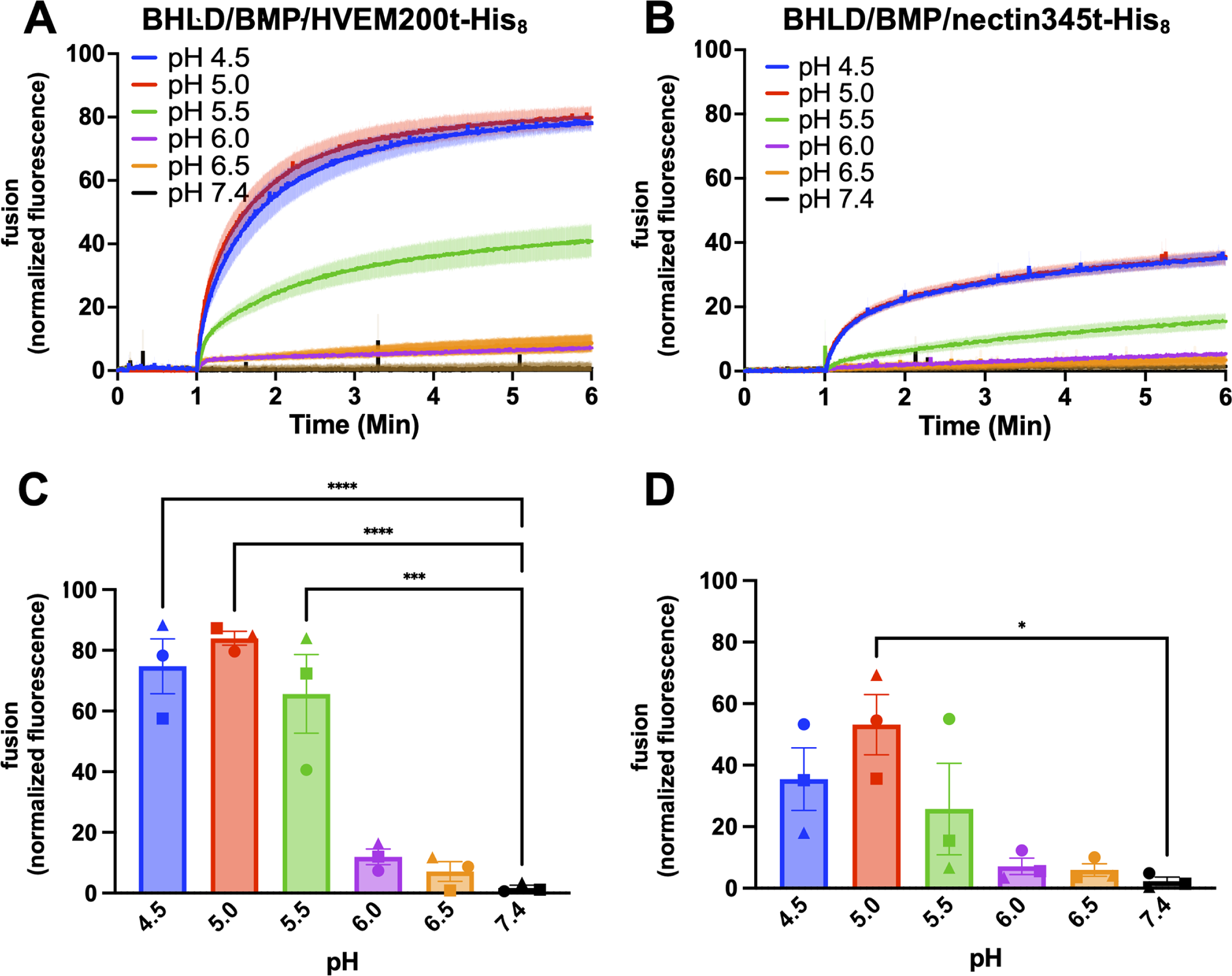
A soluble receptor (HVEM or nectin-1) in combination with low pH triggers the fusion of the VSVΔG-BHLD pseudotype *in vitro*. VSVΔG-BHLD was incubated with SRB-containing BMP LUVs at different pH in the presence of HVEM200t-His_8_ or nectin345t-His_8_. Increase in SRB fluorescence due to its dilution and dequenching reported on fusion. Lipid composition is listed in Table 1. Fluorescence was recorded and normalized to fully dequenched fluorescence by using the equation: *F_normalized_=F_t_-F_i_ / (F_max_-F_i_)^−1^ x 100%*, where *F_t_* is fluorescence at time *t, F_i_* is initial fluorescence and *F_max_* is fluorescence after the addition of Triton X-100. (A and B) For each condition, representative traces of single biological replicates, each consisting of three technical replicates, are shown. Curves represent the mean values, and the shaded area, the SEM. (C and D) The extent of fusion at 5 min post-triggering (acidification). Each data point is a biological replicate representing a mean normalized fluorescence from three technical replicates. Bars represent the mean values, and the error bars, the SEM. Each condition was done in triplicate. *: p < 0.05, **: p < 0.01, ***: p<0.001 ****: p<0.0001.

The VSVΔG-BHLD pseudotypes fused even more efficiently than HSV-1 (**Fig. 2**). For example, in the presence of HVEM200t-His_8_, the VSVΔG-BHLD pseudotypes achieved ∼80% fusion at the optimal pH of 5.0 (**Fig. 6A and C**) compared to ∼60% for HSV-1 (**Fig. 2B and E**). Moreover, the VSVΔG-BHLD pseudotypes fused more efficiently at a pH of 5.5, with ∼60% fusion (**Fig. 6AC**) compared to ∼20% for HSV-1 (**Fig. 2B and E**). Additionally, in the presence of nectin345t-His_8_, the VSVΔG-BHLD pseudotypes fused more efficiently and at a higher pH, achieving ∼50% dequenching at a pH of 5.0 (**Fig. 6B and D**) compared to ∼15% with HSV-1 (**Fig. 2C and F**). Higher errors in technical replicates (**Supplementary Fig. S4D and E**) could be due to heterogeneity in the pseudotype production. Nonetheless, we conclude that on the virion side, the essential HSV-1 entry glycoproteins gB, gD, gH, and gL are sufficient for efficient *in-vitro* fusion.

To confirm that HSV-1 entry glycoproteins were required for in-vitro fusion just as for HSV-1 entry and cell-cell fusion, we generated the VSVΔG-BD pseudotype, which lacks the gH/gL complex. The fusion levels of the VSVΔG-BD at pH 5.0 were ∼25% and ∼20% in the presence of HVEM200t-His_8_ or nectin345t-His_8_, respectively (**Supplementary Fig. S7**). These levels are much lower than those of VSVΔG-BHLD under the same conditions, ∼80% and ∼50% (**Fig. 6**), and comparable to the fusion levels of VSVΔG-BHLD in the absence of the receptor, ∼10% (**Supplementary Fig. S7**).

Fusion rates, k_fast_, were similar for HSV-1 and the VSVΔG-BHLD pseudotype regardless of the receptor, which ruled out the possibility that the differences in fusion extent between HSV-1 and the VSVΔG-BHLD pseudotype were due to the differences in fusion rates (**Supplementary Fig. S8**). This is an expected finding because fusion in both cases utilizes the same fusogen, gB, so the fusion mechanisms are expected to be similar. The differences in fusion efficiency could, then, be dictated by the amount of gB, the fusogen, on the particle surface. To test this, we directly compared the amount of gB on HSV-1 and the VSVΔG-BHLD pseudotype by Western Blot analysis. Indeed, the VSVΔG-BHLD pseudotype has more gB, on average, compared to HSV-1 (**Supplementary Fig. S9**), which could explain its higher fusion efficiency.

### In-vitro fusion of the VSVΔG-BHLD pseudotype requires a lipid with a negatively charged headgroup, cholesterol, and POPE in the target membrane

To compare the lipid requirements for the *in-vitro* fusion of VSVΔG-BHLD relative to HSV-1, we omitted single lipids (POPE, BMP, and cholesterol) from target LUVs by replacing them with the equivalent amounts of POPC (**Table 1, no POPE, no BMP, no Chol**). As with HSV-1, optimal *in-vitro* fusion with VSVΔG-BHLD required both BMP and 33% cholesterol (**Fig. 7**). However, unlike HSV-1, VSVΔG-BHLD fusion also required POPE because its omission reduced fusion ∼4-fold (**Fig. 7**). Moreover, while 17% Chol LUVs supported efficient fusion in the case of HSV-1 (**Fig. 5**), in the case of VSVΔG-BHLD, fusion in the presence of 17 and 0% cholesterol was equally low (**Fig. 7**) with a ∼4-fold reduction. Therefore, the VSVΔG-BHLD pseudotype has more stringent requirements for target membrane lipid composition than HSV-1.

**Figure 7.**
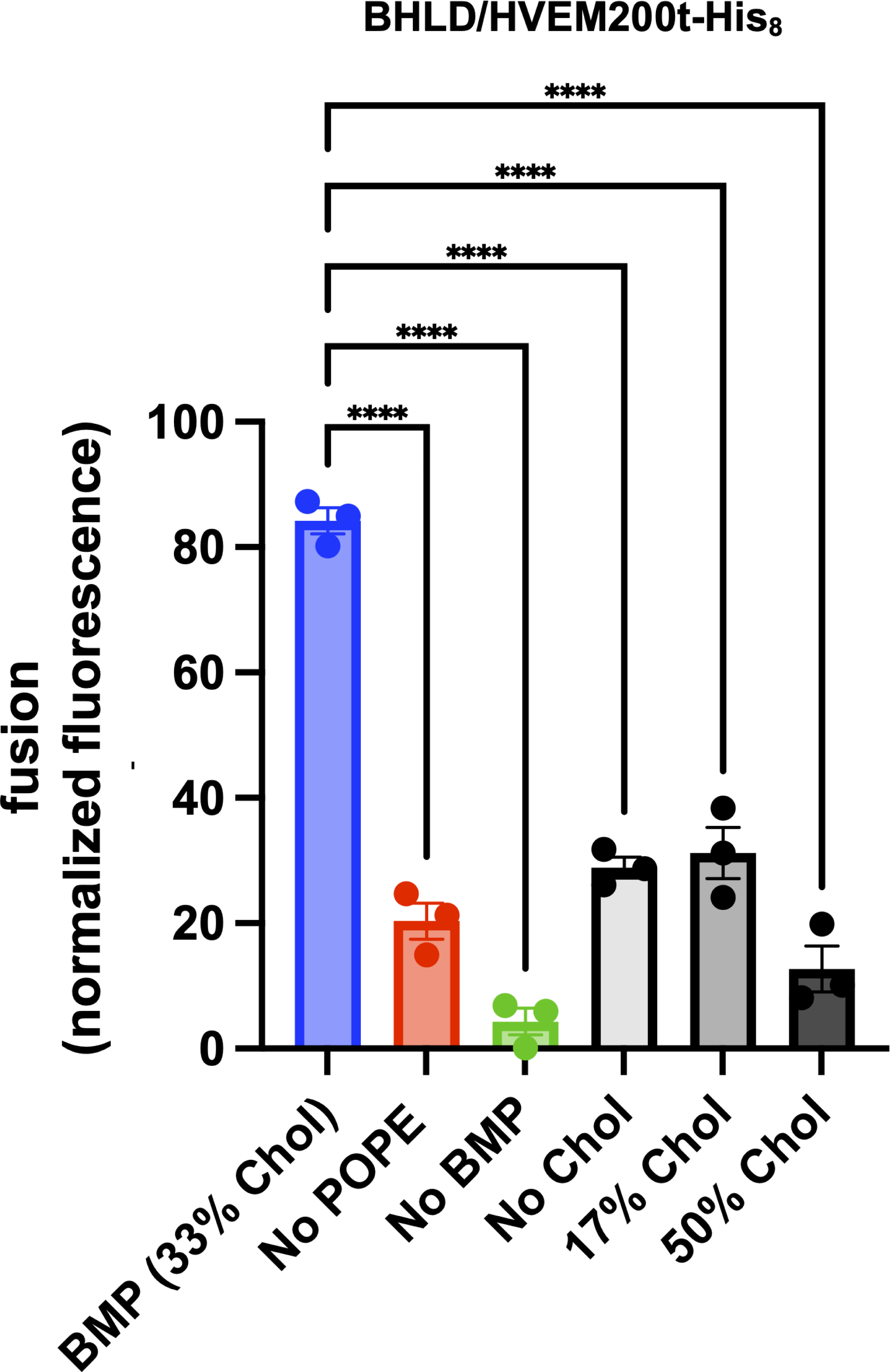
*In-vitro* fusion of the VSVΔG-BHLD pseudotype depends on the lipid composition of the target membranes. VSVΔG-BHLD was incubated with SRB-containing LUVs of different lipid compositions at pH 5.0 in the presence of HVEM200t-His_8_. Increase in SRB fluorescence due to its dilution and dequenching reported on fusion. Lipid compositions are listed in Table 1. Fluorescence was recorded at 5 min post-triggering (acidification) and normalized to fully de-quenched fluorescence by using the equation: *F_normalized_=F_t_-F_i_ / (F_max_-F_i_)^−1^ x 100%*, where *F_t_* is fluorescence at time *t, F_i_* is initial fluorescence and *F_max_* is fluorescence after the addition of Triton X-100. Each data point is a biological replicate representing a mean normalized fluorescence from three technical replicates. Each condition was done in triplicate. Bars represent the mean values, and the error bars, the SEM. Data for the BMP composition are identical to those in Figure 6. *: p < 0.05, **: p < 0.01, ***: p<0.001 ****: p<0.0001.

## DISCUSSION

### In-vitro reconstitution of HSV-1-mediated membrane fusion identifies low pH as a fusion co-trigger

HSV-1-mediated membrane fusion is a complex process, with multiple factors required on both the viral and host sides. To better define the factors that are necessary and sufficient for this process, we reconstituted membrane fusion *in vitro* with purified HSV-1 virions and synthetic target liposomes. As expected, efficient fusion required the presence of a host entry receptor, HVEM or nectin-1. These receptors were added in the form of purified ectodomains in solution and did not need to be tethered to the target membranes, which is consistent with their primary roles as fusion triggers rather than target membrane attachment factors. Importantly, we found that in addition to HVEM or nectin-1, *in-vitro* fusion also required low pH, with an optimal pH of 5.0, in contrast to cell-cell fusion, which occurs at neutral pH (28, 29).

Under *in-vitro* conditions, HVEM200t-His_8_ supported more efficient fusion than nectin345t-His_8_. Although HVEM and nectin-1 have overlapping binding sites on gD, HVEM binding requires the formation of an N-terminal hairpin in gD whereas nectin-1 binds a pre-existing site (89). Therefore, it is plausible that HVEM triggers conformational changes in gD more efficiently at low pH, at least, under *in-vitro* conditions. In contrast, nectin-1 may require additional cellular factors for efficient triggering.

Our findings are consistent with an earlier study where HSV-1 was associated with synthetic liposomes at low pH (5.0-5.3) in the presence of HVEM162t-His_6_ (equivalent to HVEM200t-His_8_ used in our study) but not nectin346t-His_6_ (90). Just as in our study, the receptors were not tethered to the liposomes. Authors proposed that the binding of soluble HVEM162t-His_6_ to gD on the surface of HSV-1 at low pH triggered a cascade of conformational changes that led to the exposure of gB fusion loops and their insertion into the liposomes (90). We hypothesize that fusion in our *in-vitro* system is triggered in much the same way.

HSV-1-mediated fusion is typically considered to be receptor-triggered rather than pH-triggered because fusion always requires an entry receptor but can occur at neutral pH. For example, HSV-1 enters some cell types, including one of its physiological targets, neurons, by fusion at the plasma membrane (19–21). Robust fusion at neutral pH is also observed in uninfected cells overexpressing the four essential HSV-1 entry glycoproteins, gB, gH, gL, and gD, so long as an entry receptor is present on target cells (28, 29). However, HSV-1 entry into many cell types, including the physiologically relevant epithelial cells and keratinocytes, occurs by endocytosis and fusion with an endosomal membrane (22-27, 30) and in some cell types, is sensitive to inhibitors of endosomal acidification (30). Given that low pH was required for fusion in our simplified system, we hypothesize that low pH is a biologically relevant co-trigger for HSV-1 fusion. The requirement for low pH as a co-trigger would explain why HSV-1 enters certain cell types by pH-dependent endocytosis. But if low pH were a fusion co-trigger, then how could fusion also occur at neutral pH? We hypothesize that in the context of cell entry at the plasma membrane or cell-cell fusion, the co-trigger role may be fulfilled by other, yet undefined, cellular factors, including proteins or other surface molecules.

### In-vitro HSV-1 fusion requirement for BMP or POPS implicates endosomal entry

On the target membrane side, efficient *in-vitro* fusion required a lipid with a negatively charged headgroup, BMP or POPS. BMP is found only in late endosomes (84, 85) and has been reported as important for entry of Dengue virus (43), VSV (46, 91), Lassa (92), and Uukuniemi virus (32). Interestingly, optimal HSV-1 fusion *in vitro* occurs at a pH of 5.0, which is typical of late endosomes. The requirement for the pH and the lipid found in late endosomes suggests that during endocytic entry into some cells, HSV-1 may fuse with the membrane of late endosomes. Further studies are needed to clarify the precise location of the HSV-1 fusion compartment across different cell types.

POPS is found in early endosomes, the inner leaflet of the plasma membrane, and viral envelopes. When present on viral envelopes, it aids cell entry [reviewed in (93)]. Interestingly, HSV-1 has been shown to activate phospholipid scramblases (94) that redistribute phosphatidylserines to the outer leaflet of the plasma membrane. Therefore, POPS could potentially play a role during HSV-1 entry by facilitating fusion at the plasma membrane or in early endosomes. POPS can also interact with calcium (95) changing the physical properties of membranes, and calcium can trigger the formation of fusion pores in artificial systems (96). Moreover, it is important for the fusion of viruses such as the Middle Eastern Respiratory Syndrome virus (97) and the Ebola virus (39). HSV-1 has also been shown to promote entry by triggering the release of intracellular calcium (98). Therefore, it is possible that under certain conditions, calcium could function as a co-trigger for HSV-1 fusion.

The third lipid with a negatively charged headgroup, POPA, did not support efficient fusion. Unlike BMP and POPS, POPA has not yet been implicated in the entry of any virus. POPA is also short-lived and is likely a messenger and metabolic precursor of other phospholipids reviewed in (99–101).

### Efficient HSV-1 fusion in vitro requires a balance of lipid order and fluidity

Efficient HSV-1 fusion *in vitro* required cholesterol, with 33% being the optimal concentration. Cholesterol modifies many of the physical properties of membranes, including fluidity, elasticity, thickness, and domain formation (71, 73, 77, 82, 102-109), reviewed in (75), in a concentration-dependent manner (71–74). Many enveloped viruses require cholesterol either in the host cell membrane or the viral envelope for entry (40, 41, 63-70). Indeed, cholesterol and other sterols, such as desmosterol, are required for efficient HSV-1 infection (57, 66, 68), for yet unclear reasons. In the case of HIV entry, cholesterol has been proposed to induce the formation of ordered lipid domains, specifically requiring the L_o_/L_d_ interface (41) and can further regulate fusion by reducing membrane lateral mobility (63). Therefore, we hypothesized that HSV-1-mediated fusion *in vitro* depended on lipid order in target membranes.

To test this, we assessed the degree of lipid order in lipid compositions used in fusion and correlated lipid order with fusion efficiency. We found that *in vitro*, HSV-1-mediated fusion levels depended on cholesterol concentration and lipid order. In the presence of 50% cholesterol, which corresponds to the high amount of ordered lipid phase, fusion levels were lowest. This suggested that high lipid order precludes HSV-1 fusion. In the absence of cholesterol, where lipids were predominantly disordered, we observed intermediate fusion levels. Efficient fusion was achieved in the presence of 17% or 33% cholesterol, which had intermediate lipid order, the latter composition being optimal for fusion. In addition to cholesterol, BMP and POPE also influenced lipid order and the opposite effects on the ordered lipid phase, with BMP decreasing it and POPE increasing it. Therefore, we conclude that efficient HSV-1-mediated fusion *in vitro* requires ordered lipid phases in target membranes and that optimal balance of lipid order and fluidity is achieved in the presence of 33% cholesterol, BMP, and POPE.

Interestingly, we found that the addition of SM to this optimal lipid composition reduced fusion levels to very low while not affecting the degree of lipid order. Therefore, ordered lipid phases formed in the presence of SM are qualitatively different from those formed in its absence and less conducive to fusion. In biological membranes, SM is typically found in ordered lipid domains, or lipid rafts (77, 78). If lipid rafts form under our experimental conditions, they appear to be, somehow, incompatible with efficient HSV-1 fusion. The optimal lipid composition for *in-vitro* HSV-1 fusion may contain very small lipid rafts. Alternatively, it may lack ordered lipid domains but, instead, have higher viscosity (83). Future studies will elucidate the nature of the ordered lipid phase formed in target LUVs and its role in HSV-1 fusion.

### The VSVΔG-BHLD pseudotypes fuse even more efficiently than HSV-1 in vitro

By reconstituting fusion with the VSVΔG-BHLD pseudotypes instead of the native HSV-1 virions, we showed that on the virus side, the four entry glycoproteins, gB, gH, gL, and gD, were sufficient for fusion. Surprisingly, the VSVΔG-BHLD pseudotypes supported higher levels of fusion *in vitro* than HSV-1. Fusion rates, k_fast_, were similar for HSV-1 and the VSVΔG-BHLD pseudotype regardless of the receptor, which suggested that the differences in fusion extent were due to the differences in fusion efficiency rather than fusion rates (**Supplementary Fig. S8**).

The reasons for this are not yet clear. However, the distinct membrane curvature of the bullet-shaped VSV pseudotypes could promote better fusion. Alternatively, the pseudotypes could have a more optimal distribution of glycoproteins on their surface. To improve the yields of the pseudotypes, we transfected 4 times more plasmid encoding gB relative to the other three plasmids. As a result, the VSVΔG-BHLD pseudotypes could contain a higher density of gB, which could potentially promote higher levels of fusion. Indeed, we found that the VSVΔG-BHLD pseudotype had, on average, more gB compared to HSV-1 (**Supplementary Fig. S9**), which could explain its higher fusion efficiency. Finally, since our pseudotypes lack any HSV-1 tegument proteins, we might not be recapitulating tegument-glycoprotein interactions (55, 110). There may be outside-in (or vice versa) signaling between the tegument proteins and membrane proteins (55, 111) that could regulate HSV-1 fusion.

Both the VSVΔG-BHLD pseudotypes and HSV-1 required cholesterol and BMP for efficient fusion. However, the pseudotype was more sensitive to cholesterol depletion and, unlike HSV-1, required POPE. POPE is known to promote the formation of the hemifusion stalk intermediate due to its negative curvature [reviewed in (101)]. If the VSVΔG-BHLD fusion depended on the formation of the hemifusion stalk as a rate-limiting step, it could be sensitive to the availability of POPE. While cholesterol is also thought to promote hemifusion by inducing negative curvature (101), it can affect membrane dynamics and fusogen activity in other ways, including but not limited to lipid ordering discussed above [reviewed in (75)], so its contributions to fusion are not limited to changing membrane curvature.

A more efficient fusion of the VSVΔG-BHLD pseudotypes *in vitro* raises the question of why these pseudotypes have a narrow tropism. In our previous work, we showed that the VSVΔG-BHLD pseudotype efficiently entered only two of the seven HSV-1 susceptible cell lines (57). Cell entry efficiency did not correlate with the type of the receptor (nectin-1 versus HVEM), their cell surface amounts, or with the route of entry (endosomal versus plasma membrane). The gB:gH:gL:gD ratios were also similar between the pseudotypes and HSV-1 (57). Earlier, we concluded that one or more of the 12 so-called non-essential HSV-1 envelope proteins, missing from the pseudotypes, could contribute to the broad tropism of HSV-1. Our work does not offer a clear explanation for the limited tropism of the pseudotypes. However, we hypothesize that it is not due to a defect in fusion but rather a defect at an earlier step in entry, e.g., endocytosis or the inability to engage a cellular factor necessary for entry at the plasma membrane.

By reconstituting HSV-1 fusion *in vitro* here, we have not only established the 4 glycoproteins and the receptor are sufficient for fusion in a minimal, bare-bones system, but have also provided evidence that low pH serves as a co-trigger of HSV-1-mediated fusion. The *in-vitro* fusion system established here can be used for a systematic exploration of the mechanism of HSV-1-mediated fusion.

## MATERIALS AND METHODS

### Cells

HEK293T (gift from John Coffin, Tufts University) and Vero (ATCC CCL-81) cells were grown in Dulbecco’s modified Eagle medium (DMEM, Lonza) containing high glucose and sodium pyruvate and supplemented with L-glutamine, 10% heat-inactivated fetal bovine serum (FBS), and 1x penicillin-streptomycin (pen-strep) solution. C10 cells (a gift from Gary Cohen, University of Pennsylvania), a clonal B78H1 derivative stably expressing human nectin-1, were grown in DMEM containing high glucose, sodium pyruvate, and L-glutamine and supplemented with 5% FBS and pen-strep solution and maintained under selection for nectin-1 expression with 250 mg/mL of G418. All mammalian cells were maintained at 37 °C and 5% CO_2_ unless otherwise noted. Sf9 cells were grown in suspension in serum-free Sf-900 II media (Gibco) containing 1x pen-strep in spinner flasks at 27 °C with aeration. For larger flasks, additional aeration was provided by an air pump through a 2-port airflow assembly.

### Plasmids

pPEP98, pPEP99, pPEP100, and pPEP101, which encode full-length gB, gD, gH, and gL from HSV-1 (strain KOS), respectively, were a gift from Patricia Spear (Northwestern University). pCMV-VSV-G encoding the full-length VSV G was a gift from Judith White (University of Virginia).

### LUV preparation and labeling

LUVs were prepared by mixing lipids in chloroform at indicated experimental compositions **(Table 1**) using the following lipids: 1-palmitoyl-2-oleoyl-sn-glycero-3-phosphocholine (POPC), 1-palmitoyl-2-oleoyl-sn-glycero-3-phospho-L-serine (POPS), 1-palmitoyl-2-oleoyl-sn-glycero-3-phosphoethanolamine (POPE), 1-palmitoyl-2-oleoyl-sn-glycero-3-phosphate (POPA), bis(monoacylglycero)phosphate (BMP), 1,2-dioleoyl-sn-glycero-3-[(N-(5-amino-1-carboxypentyl)iminodiacetic acid)succinyl] (nickel salt) DGS-NTA(Ni), Sphingomyelin (SM), or cholesterol (Chol). After being mixed, the lipids were dried under a gentle argon stream and allowed to further dry under a vacuum overnight with minimal exposure to light. The lipids were then hydrated using HBS-Citrate-SRB buffer (10 mM HEPES, 50 mM sodium citrate, 150 mM NaCl pH 7.5, 25 mM Sulforhodamine B) when used for fusion or HBS-Citrate buffer (10 mM HEPES, 50 mM sodium citrate, 150 mM NaCl pH 7.5) when used for Laurdan fluorescence spectroscopy. The lipids were warmed to 37 °C and vortexed several times before being first frozen in liquid nitrogen and then thawed in a water bath at 37°C. The lipid suspension went through 5 freeze/thaw cycles before being extruded through a 100-nm filter 21 times. To remove excess SRB, LUVs using a Sephadex G-25 column equilibrated with HBS-Citrate buffer.

### LUV Quantification

Prior to bulk fusion experiments, all LUV preparations were quantified and measured using multi-angle light scattering (MALS). MALS analysis (Dawn Heleos II, Wyatt Astra 7.3) included measuring the radius of gyration (r_g_) and concentration of LUVs, which was then compared to theoretical concentrations calculated as described in (112). An additional detector for dynamic light scattering (DLS) was used to report the hydrodynamic radius (r_h_) of LUVs to quantify size distribution. LUV extrusions were measured in batch mode using a microcuvette with a sample volume of 20 μL. Dilutions of LUVs were prepared, and appropriately diluted to keep the sample within the dynamic range of the instrument. Light scattering (LS) detection limits were previously determined and validated using NIST standard polystyrene beads of 100-nm and 200-nm diameter. Representative measurements include the LUV concentration (**Supplementary Figure S1A**), distribution (r_h_) (**Supplementary Figure S1B**), and radius size (r_g_) (**Supplementary Figure S1C**).

Potential dequenching (dilution of SRB) is dependent on the relative increase in volume after fusion. As such, LUV volume is sensitive to changes in radius size during the extrusion process, which can influence dequenching potential. Therefore, consistent results are dependent on consistent radii. The radii were calculated using the Lorenz-Mie coated sphere model and the following parameters: shell thickness of 5 nm (representing typical bilayer thickness) and refractive indices of 1.46 and 1.33 for the shell (lipid bilayer) and the sphere (water), respectively. R_g_, r_h,_ and model fit were all used to determine the quality of lipid extrusions. The LUVs extruded through a 100-nm filter had r_g_ values of 50-60 ± 1-3 nm. Inconsistent bulk fusion traces could be traced back to LUV preparations where the measured radii had error values ±10-25 nm and wider r_h_ distribution, in addition to deviation from the Lorenz-Mie coated sphere model, indicative of insufficient extrusion and the presence of multilamellar vesicles.

### Laurdan Fluorescence Spectroscopy

LUVs were prepared in the same manner as LUVs used for fusion except that the LUVs contained 1% Laurdan, were rehydrated in HBS-Citrate without SRB, and did not require desalting. Fluorescence spectra were recorded on a Horiba Quantamaster 8000 equipped with a Peltier heated cuvette holder with stirring capability. The holder had constant stirring at 700 rpm and variable temperature. Fluorescence excitation and emission wavelengths were 350 nm and 400-600 nm, with a slit width of 5 nm for excitation and 1 nm for emission. Approximately 1.5 mg/mL of LUVs were used to obtain emissions scans. The temperature was ramped up automatically, and once the cuvette reached the target temperature, the emission scan was repeated. Raw fluorescence values were used to calculate GP values using the equation GP = (I_440nm_-I_480(494)nm_)/(I_440nm_+I_480(494)nm_)^−1^.

### HSV-1 propagation and purification

HSV-1 strain GS3217 (an F strain derivative encoding an NLS-tdTomato transgene expressed from an IE promoter, a gift from Greg Smith, Northwestern University) was propagated and titrated on Vero cells (ATCC). To produce large viral stocks, approximately 2×10^9^ adherent Vero cells (10 T175 flasks, 2×10^7^ cells/flask) were infected with HSV-1 at MOI 0.01 by replacing cell media with virus diluted in minimal media Opti-MEM (ThermoFisher). The virus was allowed to adhere to cells at 37 °C in 5% CO_2_ for 1 hour with minimal agitation every 15 minutes. After 1 hour, the virus-containing media was removed and replaced with 10% FBS DMEM. Cells were incubated at 37 °C in 5% CO_2_, and the supernatant was collected 48-72 hours later.

To ensure batch-to-batch consistency, HSV-1 virions were purified by density gradient centrifugation using a 10-50% iodixanol gradient to separate them from the L-particles, according to published protocols (113, 114) **(Supplementary Figure S1D**). Briefly, virus-containing media was clarified of cell debris by centrifugation at 1,500 *g* for 15 minutes, twice. The virus was then pelleted in an SW27 rotor at 28,000 *g* for 90 min. The supernatant media was removed and replaced with a minimal volume of HNE buffer (50 mM HEPES, 150 mM NaCl, pH 7.4) to resuspend the viral pellets to a combined volume of approximately 1 mL. The viral pellets were first agitated and then incubated in HNE buffer overnight at 4 °C. Iodixanol step gradients from 10-50% in 10% increments were prepared and allowed to equilibrate to form a continuous gradient overnight at 4 °C. The viral pellets were resuspended HNE buffer by pipetting gently and pooled together. The pooled virus was then loaded onto the step gradients and density purified using an SW41 rotor at 160,000 *g* for 4 hours, with a “no brake” stopping setting. The resulting viral band was collected using a side-puncture technique, aliquoted, and stored at −80 °C. Viral titers were determined by plaque assay in Vero cells as described in (113, 114).

The purified HSV-1 virions were then analyzed using MALS and DLS in batch mode using a Wyatt Dawn Heleos II in batch as described in the LUV quantification section with some modifications. The radii were calculated using the Lorenz Mie homogenous sphere light scattering model and Rayleigh-Gans Sphere model using a refractive index value of 1.41 (115) and were on average ∼98 nm (representative data shown in **Supplementary Figure S1G**). Although both models had their caveats, e.g., Rayleigh-Gans is more accurate when the refractive index of the sample is close to the environment and Lorenz-Mie assumes the particle is homogenous, the experimentally determined radii agreed with the literature values of 92-115 nm (54, 55). Concentration values were between 1×10^10^ and 1×10^11^ particles per mL, on average (representative data shown in **Supplementary Figure S1E**). Using the titers and the particle counts, we calculated the particle-per-PFU ratios. Most HSV-1 preparations had a particle-to-PFU ratio of 20-30, on average (representative data shown in **Supplementary Figure S1H**). These values are close to the initially reported value of 10 (116) and the more recently reported values of 20-23 (56), which are used as gold standards of HSV-1 virion quality. Only virion preparations with a particle-to-PFU ratio of 20-30 were used in our experiments because values of >30 produced inconsistent results. A representative size distribution as measured by DLS (**Supplementary Figure S1F**) shows a typical preparation. Wider distributions were often seen with preparations with particle-to-PFU ratios of >30 and were used for initial quality assessment.

### VSVΔG-BHLD and VSVΔG-BD Pseudotype Generation and Purification

The VSVΔG-BHLD pseudotype was prepared as described in (88) with minor modifications. Briefly, HEK293T cells (1.7 x 10^7^ cells/T175 flask) were transfected with 7.5 μg each of pPEP99, pPEP100, pPEP101, and 30 μg of pPEP98 with polyethyleneimine (PEI: 1mg/mL) at a 3:1 ratio. To generate the VSVΔG-BD pseudotype, pPEP100 and pPEP101 were replaced with the pCAGGS vector. Ten T175 flasks were used for producing enough purified virus for bulk fusion. After 24 hours post-transfection at 37 °C, cells were infected with the VSVΔG-G “helper” pseudotype at an MOI of 3 at 30 °C. 48 hours post-infection, the supernatant was collected, and the virus was purified by density centrifugation. The initial purification step was to clear cell debris by centrifuging the sample through a 12% iodixanol cushion at 1,500 g for 15 minutes. The virus was then pelleted using an SW27 rotor at 28,000 *g* for 90 min. The supernatant was removed and replaced with a minimal volume of HNE buffer (50 mM HEPES, 150 mM NaCl, pH 7.4) to resuspend the viral pellets in a combined volume of approximately 1 mL. The viral pellets were agitated and then incubated in HNE buffer overnight at 4 °C, undisturbed. Iodixanol step gradients from 15-35% in 5% increments were prepared and allowed to equilibrate to a continuous gradient overnight at 4 °C. The viral pellets were resuspended in HNE buffer by gentle pipetting and pooled together. The pooled virus was then loaded onto the continuous gradients and density purified using an SW41 rotor at 160,000 *g* for 90 min, with a “no brake” stopping setting. The resulting viral band was collected using a side-puncture technique, aliquoted, and stored at −80 °C.

Viral titers (Infectious Units or IU) on C10 cells were calculated from the number of GFP-positive cells per inoculum volume. The purified virus was also characterized and counted using MALS and DLS using a Wyatt Dawn Heleos II in batch mode as described in the LUV quantification section. However, due to the shape of the virus interpretation using software (Astra 7.3) had to be modified. The bullet shape could be approximated to a rod and the best approximation to obtain particle number and length (as opposed to radius) requires a user-defined cylinder radius and refractive index (Rayleight-Gans: Rod). We used a radius of 35 nm that we determined from negative stain electron microscopy of purified BHLD particles. As there are no reported refractive index values for VSV or any other *Rhabdoviridae*, we used a value of 1.55, which approximates an extracellular vesicle containing 1/3 each protein (1.584) 1/3 lipid (1.460) and 1/3 nucleic acid (1.620).

### Expression and Purification of HVEM and Nectin-1 ectodomains

Genes encoding His-tagged ectodomain versions of the HSV-1 gD receptors HVEM (HVEM200t) or nectin-1 (nectin345t) were synthesized by GenScript (GenBank accession numbers AF060231 and U70321, respectively). Native signal peptide sequences were replaced with those of honeybee melittin and terminated with 8 histidines. The resulting constructs HVEM200t-His_8_ and nectin345t-His_8_ are nearly identical to ones previously characterized in the literature (90, 117, 118). HVEM200t-His_8_ used here is identical to HVEM162t-His_6_ reported elsewhere (118) but uses a different numbering convention (residue 1 corresponds to the first methionine rather than the first residue of the mature protein, after the cleavage of the signal sequence). The sequences were subcloned into the baculovirus expression vector pFastBac1 (Invitrogen). The recombinant, soluble His-tagged versions of the receptors were produced in Sf9 cells infected with recombinant baculoviruses as described below.

Recombinant bacmids and recombinant baculovirus stocks were produced by GenScript. The high-titer P2 was used to make P3 baculovirus for a large-scale expression. 4 L of Sf9 cells grown in suspension were infected with P3 baculovirus at a ratio of 1:250. At 48 hours post-infection, the cell suspension was centrifuged at 4,500 *g* for 30 min at 4 °C and the pellet was discarded. The cell supernatant was then filtered through a 0.22-μm pore size filter. The filtered supernatant was then loaded onto a 5-mL HisTrap Excel column (Cytiva) at a rate of 5 mL/min. The column was then washed with 10 column volumes each of the following concentrations of imidazole: 0 mM, 10 mM, 25 mM, and 50 mM in 150 mM PBS, pH 7.4 buffer. Protein was eluted with 250 mM imidazole in 150 mM PBS, pH 7.4 buffer. The eluate was further purified using size-exclusion chromatography (Superose 6 column, Cytiva) to remove aggregates and remaining contaminants. Peak fractions were pooled, concentrated to 5 mg/mL, and stored at −80 °C. Representative Coomassie-stained gels of purified proteins are shown in **Supplementary Figures S3A and B.**

### In-vitro Bulk Fusion Assay with HSV-1 and the VSVΔG-BHLD pseudotype

Approximately 2-5×10^7^ PFU (∼0.4-1×10^9^ particles) of HSV-1 or ∼0.4-1×10^9^ particles of the VSVΔG-BHLD pseudotype were mixed with 0.4-1 μM LUVs (0.5-1×10^9^ particles) with or without 0.8 μM HVEM200t-His_8_ or nectin345t-His_8_ in HBS-Citrate buffer in a quartz cuvette to a total volume of 2 mL. Fluorescence was recorded using a Horiba Quantamaster 8000 equipped with a Peltier heated cuvette holder with stirring capability. The holder was kept at 37 °C with constant stirring at 700 rpm. Fluorescence excitation and emission wavelengths were set to 565 nm and 586 nm with a slit width of 5 nm. The sample was incubated in the cuvette at 37 °C with stirring for 10 min before data collection. Fluorescence was then recorded for 1 min to obtain baseline fluorescence. At 1 min, pH was reduced using predetermined amounts of 1 M HCl, or the soluble receptor was added. After 5 min, post pH shift or receptor addition, the sample was fully de-quenched by the addition of 10% Triton X-100 to a final concentration of 0.1%. Fully de-quenched fluorescence was recorded for 30 s and used for normalizing the fluorescence data. Data were normalized using the equation F_normalized_=F_t_-F_i_ / (F_max_-F_i_)^−1^, where F_i_ is the initial, baseline fluorescence, F_t_ is the fluorescence at time *t*, and F_max_ is fluorescence after the addition of Triton X-100. Normalized values of the technical replicates were then averaged. Fluorescence at 5 min post triggering, pH shift, or receptor addition, were considered endpoint values for comparison. Where applicable, the normalized data were also fit using the following two-phase (exponential) association analysis built into GraphPad PRISM9, Y = Y_0_ + SpanFast x (1-e ^(-K^ ^(t)^) + SpanSlow x (1-e^(-K^ ^(t))^), where SpanFast = (Plateau - Y) x PercentFast x 0.01 and SpanSlow = (Plateau - Y_0_) x (100-PercentFast) x 0.01.

### HSV-1 Inactivation Assay

Approximately 5×10^5^ PFUs of HSV-1 in 1mL of HBS-Citrate buffer were incubated at 37 °C or 4 °C with or without 1 μM HVEM200t-His_8_ for 1 hour. pH was then shifted to the indicated pH using predetermined amounts of 1 M HCl and incubated at 37 °C (or held at 4 °C for the pH 5.0/4 °C condition). After 1 hour, the sample was back titrated to pH 7.4 using 1 M NaOH. For neutral pH conditions, the buffer was used instead of HCl or NaOH. The sample was then kept at 4 °C before being diluted in Optimem and immediately used for infection. Approximately 50 PFUs were used to infect 1.5×10^5^ Vero cells per well, in duplicate, in a 6-well plate. The inoculum was replaced with 10% FBS DMEM containing 1.25% carboxymethylcellulose (CMC) after 1 hour. Plaques were grown for 48-72 hours, at which point media was removed and cells were stained using 1% crystal violet dissolved in 50% methanol. Cells were destained using deionized water, and plaques were counted, with counts averaged between the two wells. Plaques were normalized to the control condition of no receptor at pH 7.4.

### Western Blotting

Purified, concentrated HSV-1 or VSVDG-BHLD virions (3 x 10^7^ particles) were mixed with SDS-PAGE loading dye and boiled for 15 min. Samples were loaded onto 4-15% precast polyacrylamide gels (Bio-Rad), separated by SDS-PAGE, and transferred onto a nitrocellulose membrane. The membrane was blocked with TBS buffer containing 0.05% Tween (TBST) and 5% milk for 1 hour at room temperature. The membrane was incubated with the polyclonal rabbit anti-HSV-1-gB antibody R68 at a 1:5000 dilution in TBST at 4 °C overnight. The membrane was washed and incubated with goat anti-rabbit LI-COR IR800 secondary antibody at a 1:5000 dilution in TBST for 1 hour at room temperature. After washing, the membrane was imaged using a LI-COR Odyssey imager. R68 pAb was a gift from G. H. Cohen and R. J. Eisenberg (U. Pennsylvania).

### Statistics

Statistical analysis was done for each experiment using one-way ANOVA with multiple-comparison Tukey adjustment as implemented in GraphPad PRISM 9.

## ACKNOWLEDGEMENTS

We thank Gary Cohen (University of Pennsylvania) for the gift of C10 cells; Patricia Spear, (Northwestern University) for the gift of HSV-1 glycoprotein plasmids; John Coffin (Tufts University) for the gift of HEK293T cells; Gregory Smith (Northwestern University) for the gift of HSV-1 GS3217 strain; and Michael Whitt (University of Tennessee) and Judith White (University of Virginia) for the gift of the VSVΔG-GFP pseudotyping platform. We thank James Munro (University of Massachusetts), for help with the in-vitro fusion assay. We thank John Coffin (Tufts University), Karl Munger (Tufts University), Yu-Shan Lin (Tufts University), and Susan Daniel (Cornell University) for helpful discussions.

Research reported in this publication was supported by the National Institutes of Health under award numbers T32GM731042 (J.M.R.), T32GM731043 (J.M.R.), R21AI160821 (E.E.H.), and R21AI145272 (E.E.H.) from the National Institutes of Health, and by a Faculty Scholar grant 55108533 from Howard Hughes Medical Institute (E.E.H.). The content is solely the responsibility of the authors and does not necessarily represent the official views of the National Institutes of Health.

J.M.R. designed the experiments, conducted the experiments, analyzed the data, and generated hypotheses. A.C.-Z. conducted experiments analyzed the data, and generated hypotheses. A.B. conducted experiments, analyzed the data, and generated hypotheses. E.E.H. designed experiments, analyzed the data, and generated hypotheses. All authors wrote the manuscript.

We dedicate this publication to the memory of Roselyn J. Eisenberg (University of Pennsylvania).

